# DSCAM preserves neuronal order during radial migration by antagonizing N-Cadherin adhesion and UNC5c repulsion

**DOI:** 10.1101/2025.06.16.659927

**Authors:** Tao Yang, Ty Hergenreder, Macy W. Veling, Wei-Chih Chang, Elaine J. Cui, Yu Wang, Peter G. Fuerst, Roman J. Giger, Xiao-Feng Zhao

**Affiliations:** Department of Neurology, University of Michigan, Ann Arbor, MI 48109, USA; Department of Cell and Developmental Biology, University of Michigan, Ann Arbor, MI 48109, USA; Life Sciences Institute, University of Michigan, Ann Arbor, MI 48109, USA; Molecular and Integrative Physiology, University of Michigan, Ann Arbor, MI 48109, USA; Wake Forest University, Winston-Salem, NC 28304, USA

## Abstract

Proper radial migration of developing cortical neurons is critical for the “inside-out” laminar organization of the mammalian cerebral cortex. While substantial progress has been made in our understanding of how neurons terminate migration at the cortical plate (CP)/marginal layer (ML) boundary, the mechanisms that control the sequential order of migrating neurons remain poorly understood. Here, we show that radial migration of upper-layer pyramidal neurons is transiently restrained within the upper cortical plate (UCP), where migrating neurons assemble into vertically organized radial queues that maintain inter-neuronal spacing and migratory order. Using time-lapse imaging of embryonic mouse cortex, we found that leading neurons transiently delay nucleokinesis of trailing neurons, thereby synchronizing neuronal movement within the UCP. Loss of Down syndrome cell adhesion molecule (DSCAM) disrupts this coordinated migratory behavior, allowing trailing neurons to bypass preceding neurons, resulting in abnormal neuronal accumulation near the CP/ML boundary. Mechanistically, DSCAM localizes to membrane interfaces between adjacent migrating neurons. At these sites, DSCAM suppresses N-cadherin-mediated intercellular adhesion, limits F-actin assembly, and inhibits expansion of the proximal cytoplasmic dilation required for nucleokinesis. *Dscam*-deficient neurons exhibit increased cadherin puncta, more neuron-neuron contacts, and accelerated migration through the UCP. Rescue experiments demonstrate that expressing the extracellular domain of *Dscam* is sufficient to restore ordered migration, suggesting that an extracellular mechanism regulates orderly neuronal migration. In addition, we identify UNC5c-mediated signaling as a counterbalancing force that promotes upward migration within the UCP. Together, our findings identify radial neuronal queues as a previously unrecognized intermediate structure in cortical morphogenesis and establish DSCAM-dependent regulation of inter-neuronal spacing as a mechanism that preserves the migratory order during inside-out corticogenesis.

## INTRODUCTION

The development of the cerebral cortex occurs in an “inside-out” fashion (also referred to as inside-out neurogenesis), a defining feature of cortical development in mammals^1^. During cortico-genesis, pyramidal neurons are generated sequentially, with early-born neurons forming the inner cortical layers, while later-born neurons migrate past them to establish new layers atop the existing ones^2–4^. Although this laminar organization is well described, the direct contribution of radially migrating neurons to cortical morphogenesis remains poorly understood. In several neurodevelopmental disorders, including Down syndrome^5,6^ and Autism spectrum disorder^7,8^, disruptions in neuronal migration and cortical organization can impair proper cortical assembly. Therefore, understanding how radially migrating neurons contribute to corticogenesis is fundamental for elucidating mechanisms underlying these disorders.

In mammals, pyramidal neurons are generated in the ventricular zone (VZ) and subventricular zone (SVZ) of the embryonic dorsal telencephalon^3,9^. Newly born pyramidal neurons undergo radial migration to generate cortical layers^10^. During this process, later-born pyramidal neurons migrate past earlier-born neurons and settle in more superficial positions within the cortical plate (CP), giving rise to the “inside-out” pattern of cortical layer formation. During radial migration, neurons extend a leading process toward the pial surface, and the soma advances through saltatory nucleokinesis^11^. Lamellipodia assemble at the proximal end of the leading process, expanding the portion of the leading process into a cytoplasmic dilation^12^. This newly formed dilation facilitates rapid somal translocation through the proximal cytoplasmic dilation of the leading process (PCDLP)^13^, known as a nucleokinesis^14^. Despite extensive progress in understanding cortical development, the molecular mechanisms that determine when and where migrating neurons bypass preceding neurons and terminate migration at their appropriate position remain incompletely understood.

Research on Reelin and DSCAM sheds light on how radial migration contributes to cortical development. Reelin, secreted by Cajal-Retzius cells in the molecular layer (ML), decreases the adhesion between nascent pyramidal neuron and radial glia, facilitating their detachment^15^. Once detached, migrating neurons stop maintaining their bipolar morphology and instead exhibit dynamic behavior, extending and retracting multiple highly mobile processes repeatedly^16^. The location of Reelin expression ensures that new neurons reach the top of CP before detaching from radial glia. DSCAM has been implicated in regulating neuronal positioning within the developing cortex. Specifically, it has been shown that DSCAM is required for migrating nascent neurons to bypass their post-migratory predecessor and migrate across the CP to expand the cortical layer, thereby enabling cortico-genesis in an “inside-out” manner^16^. However, this process requires that migrating neurons maintain their sequential order during their radial migration, a feature that has not been directly examined. Maintaining this sequential order may be particularly important, given that the morphogenesis of the upper cortical layers (layer II-IV) undergoes a prolonged maturation phase. In mice, neurons of upper cortical layers (UCL) are generated between embryonic day (E) 14.5 and E16.5 and migrate into the CP within two days. However, the UCL morphogenesis continues from E19 through postnatal day 7^17^, during which radial-migrating neurons accumulate in the developing CP. Despite recent research advancing our understanding of this process, much remains unknown about how nascent pyramidal neurons behave during this waiting period.

DSCAM is a type I transmembrane cell adhesion molecule of the immunoglobulin superfamily^18^. Its name originates from its location within the Down syndrome critical region of human Chromosome 21^19–23^. The fly homolog Dscam1 exhibits extraordinary isoform diversity of the ectodomain, mediating iso-neuronal self-avoidance and tiling, preventing excessive overlap of dendrites or axons^24–29^. Although mouse DSCAM lacks this splice diversity, it nevertheless mediates both iso- and hetero-neuronal avoidance^30^. DSCAM’s roles in vertebrate neurodevelopment have been demonstrated by a series of studies in the mouse retina. In the absence of DSCAM, the retinal mosaic becomes disorganized, neuronal somata cluster together, and neurites fasciculate together ^31–33^. Outside the retina, DSCAM facilitates UCL morphogenesis. DSCAM is necessary to guide newly arriving neurons past their post-migratory leading neuron predecessors, entering the emerging cortical layer^16^. Thus, DSCAM directly regulates cortical thickness. DSCAM acts as a critical regulator of an adhesion code in both retina and cortex, in part by masking cadherin-mediated adhesion ^16,34^. DSCAM-mediated adhesion regulation is also observed in the midbrain, where DSCAM facilitates nascent neuron detachment from the VZ. Without DSCAM, blockade of Rap1 and N-cadherin localization, neurons remain attached to the VZ and are unable to initiate radial migration^35^. The cell autonomy of DSCAM in the developing cortex, how it masks cadherin cell adhesion, and whether the DSCAM cytoplasmic domain is involved have not yet been explored.

Here, we report that radial migration is largely suspended in the UCP of wild-type (WT) mice, whereas nascent neurons continue migrating across the UCP in *Dscam* null mutants. We observed that migrating WT neurons assemble into vertically aligned multicellular structures, called radial queues, within the UCP. In these queues, leading neurons temporarily block the radial migration of the following neurons, thereby preserving neuronal spacing and migratory order. We observed that DSCAM is localized to the plasma membrane between adjacent neurons and coincides with a paucity of cadherin-mediated adhesions. We show that neuronal DSCAM is required to maintain orderly migration within radial queues, consistent with a model in which DSCAM restricts expansion of the proximal cytoplasmic dilation of the leading process (PCDLP) in trailing neurons. In the absence of DSCAM, migrating neurons form excessive cadherin-mediated adhesions, promoting PCDLP expansion and nucleokinesis. We further identify UNC5c-mediated repulsion as a counterbalancing force that promotes upward migration within the UCP. Together, our findings suggest that migratory order is actively maintained before terminal detachment at the CP/ML boundary, revealing a mechanism that preserves the sequential organization required for inside-out corticogenesis.

## RESULTS

### DSCAM regulates the distribution of radially migrating neurons in the upper cortical plate

To better understand neuronal behavior during radial migration, we labeled migrating neurons using *in-utero* electroporation (IUE; see details in Methods). Plasmids encoding fluorescent proteins were injected into the lateral ventricle of E14.5 WT or *Dscam^2j/2j^* (a frameshift null-allele mutation of *Dscam*^33^, hereafter referred to as *Dscam^-/-^*) embryos and electroporated. At E18.5, the offspring of the dorsal neural progenitor cells (NPCs) have differentiated into nascent neurons. Many of these neurons migrate radially to the upper half of the cortical plate (UCP), while the rest remain in the lower half (LCP). As expected, in both WT and *Dscam^-/-^* embryos, the radial-migrating neurons exhibited typical bipolar morphology, characterized by a leading process, cell body, and trailing process (Fig. 1a, b, d, and e). In WT neocortex, labeled migrating neurons distribute evenly across the UCP (Fig. 1a, WT). The detailed images (Fig. 1b) show that multiple neurons (arrowheads) migrated along the same axis in WT. These neurons’ cell bodies were spaced apart to form a broadly distributed queue. In *Dscam^-/-^* neocortex, migrating neurons accumulated below the CP/ML border (Fig. 1a, *Dscam^-/-^*; 1b, *Dscam^-/-^*, for detail). To quantify the distribution of migrating neurons in the UCP, we measured the distance of individual neurons to the CP/ML border (Fig. 1c). In WT neocortex, neurons were distributed in a wide area with an average distance of -52.1±30.2 µm below the CP/ML border. In *Dscam^-/-^* littermates, neurons accumulated in a narrow space, with an average distance of -36.0±26.2 µm below the CP/ML border (Fig. 1c).

**Fig. 1.**
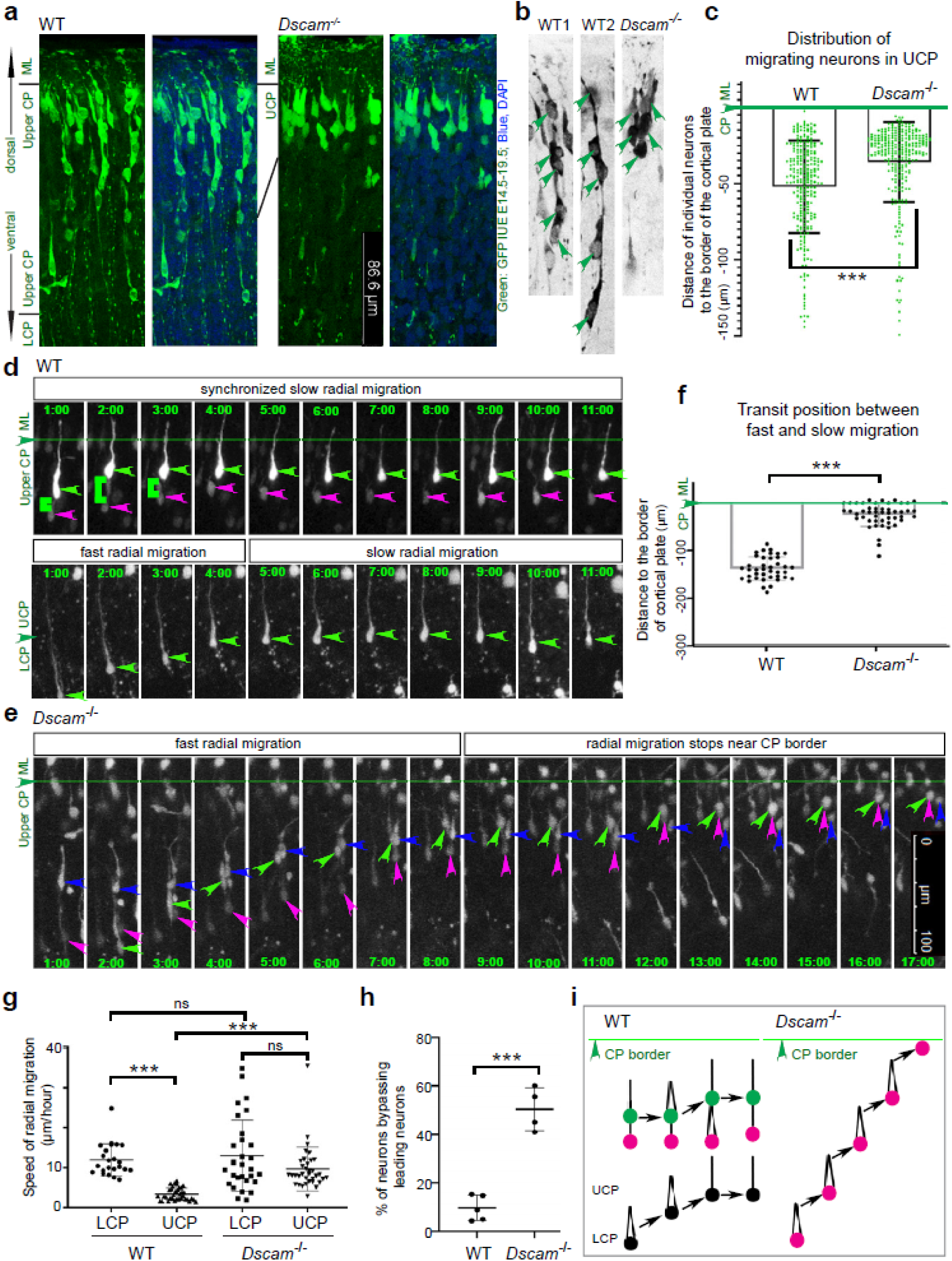
Synchronized radial migration in upper cortical plate (UCP) is disrupted in *Dscam^-/-^*mutant. **a** Distribution of GFP^+^ pyramidal neurons (green) in embryonic day (E) 18.5 UCP with DAPI counterstaining (blue) following *in-utero electroporation* (IUE) at E14.5. WT neurons are distributed in a wide area in the UCP, while *Dscam^-/-^* neurons accumulate in a narrow band at the CP/ML border. **b** In WT1 and WT2, higher magnification images show multiple GFP^+^ pyramidal neurons (green arrowheads labeled) distributed evenly along the same radial axis. In *Dscam^-/-^*, GFP^+^ pyramidal neurons cluster at the dorsal border of the UCP. **c** Quantification of distance of individual GFP^+^ neurons from the CP/ML border in 3 WT and 4 *Dscam^-/-^* brains. Mann-Whitney U test. **d,e** Time-lapse imaging shows the behavior of radial-migrating neurons in the CP: (**d**) WT top: synchronized radial migration in the UCP. Green arrowheads label the leading migrating neuron. Magenta arrowheads label the trailing neuron. Green brackets label the proximal cytoplasmic dilation of leading process (PCDLP). WT bottom: Fast radial migration is suspended when the neuron (Green arrowheads) enters the UCP. (**e**) In *Dscam^-/-^* cortex, fast radial migration occurs across the cortical plate. Green line and arrowhead label the CP/ML border. **f** Quantification of transit position of migrating neurons relative to the CP/ML border. Data points represent individual migrating neurons from 4 WT and 5 *Dscam^-/-^* brains. Mann-Whitney U test. **g** Quantification of neuron migratory velocity in the LCP and UCP from 4 WT and 4 *Dscam^-/-^* brains. Kruskal-Wallis test. **h** Quantification of the % of neurons bypassing their corresponding leading neuron. Data points represent individual animals. Student’s t-test. ns, not significant, and ***p<0.001. **i** Schematic of synchronized radial migration in the UCP. WT neurons in a queue, color-coded black (most apical, last born), pink (intermediate), and green (first born, most basal), maintain a sequence. In the *Dscam^-/-^* brain, there is no queuing, and a given neuron migrates rapidly in the basal direction.

### DSCAM is required for synchronized radial migration in the upper cortical plate

Time-lapse imaging was performed to monitor radial migration behavior of E18.5 WT and *Dscam^-/-^* neurons labeled by IUE at E14.5. In WT neocortex, radial migration mostly stalls in the UCP (Fig. 1d, upper panel). In contrast, neurons migrated rapidly in the LCP but slowed dramatically upon entering the UCP (Fig. 1d, lower panel; 1-4 hours shows the fast migration; 4-11 hours shows the stalled migration). The average migratory speed in the LCP was 12.0±4.0 µm/hour, while in the UCP, speed decreased to 3.4±1.6 µm/hour (Fig. 1g). Tracing individual neurons revealed that high-speed radial migration was halted at 137.7±24.4 µm below the CP/ML boundary, corresponding to the transition into the UCP (Fig. 1d, lower panel and 1f). In *Dscam^-/-^* neocortex, neurons migrated rapidly across both LCP and UCP, with no noticeable reduction in speed (Fig. 1e and 1f). In *Dscam^-/-^*, the speed of migration was 13.0±8.9 µm/hour in LCP, and 9.7±5.5 µm/hour in UCP. The radial migration speed in *Dscam^-/-^* UCP is comparable to that in *Dscam^-/-^*LCP and WT LCP (Fig. 1g). Radial migration speed in UCP was much slower in WT than in *Dscam^-/-^*. These data indicate that DSCAM suspends rapid radial migration in the UCP.

Among radial-migrating neurons in UCP, the leading neurons appear to set the pace for the trailing neurons (Fig. 1d, upper panel). The leading neuron (green arrowheads) moved 14.9 µm dorsally (Fig. 1d, upper panel, 1∼2 hours). Then, the following neuron (magenta arrowheads) immediately moved 14.3 µm to occupy the space vacated by the leading neuron (Fig. 1d, upper panel, 2∼4 hours). As a result, the leading neurons and following neurons maintained their sequential order. Notably, they kept their distance during their radial migration in the UCP. This observation demonstrates a synchronized radial migration in the UCP. In *Dscam^-/-^* neocortex, migrating neurons failed to maintain their synchronized movement (Fig. 1e). Time-lapse recordings revealed that the following neuron (green arrowheads) bypassed one leading neuron (magenta arrowheads, Fig. 1e, 2-4 hours) and then another (blue arrowheads, Fig. 1e, 4-8 hours). Interestingly, the newly formed following neuron (magenta) also bypassed its leading neuron later (blue arrowheads, Fig. 1e, 9-12 hours). Finally, all these neurons ceased migration just below the dorsal border of the CP (Fig. 1e, 13-17 hours). These data suggest a crucial role for DSCAM in maintaining synchronized radial migration with fixed sequential order and distance (Fig. 1i).

We quantified the synchronized radial migration in leading and trailing neuron pairs. In WT neocortex, 9.7±5.2% of the following neurons bypassed their leading neurons during the time-lapse recording. In *Dscam^-/-^*neocortex, the number increased to 50.4±8.9% (Fig. 1h). This finding supports a model in which DSCAM is required to maintain synchronized migration, preserving both sequential order and spacing among migrating neurons in the UCP (Fig. 1i).

### *Dscam* expression marks aligned column of migrating neurons in the upper cortical plate

Our previous work showed that *Dscam* expression increased dramatically in the UCP at E19^16^. Whether this pattern correlates with synchronized radial migration remains unclear. *Dscam in-situ* hybridization (ISH) of E19 neocortex showed that *Dscam* expressing cells are organized into vertical columns covering the UCP (Suppl. Fig. 1a), from the top to the base of the UCP (Suppl. Fig. 1a and 1c, square bracket labeled), with an average length of 148.2±26.5 µm (Suppl. Fig. 1b). In contrast, these *Dscam*^+^ columns in *Dscam^-/-^* brain were diminished (Suppl. Fig. 1a). By labeling migrating neurons with RFP via IUE at E14.5, we found that radial migratory RFP^+^ neurons expressed *Dscam* at E19 (Suppl. Fig. 1c and 1d, green arrowheads) in both the UCP and to a lesser degree in the LCP. The higher speed of radial migration in the LCP indicates that *Dscam* expression alone does not impede migration of a single, non-contacting cell. In contrast, migrating neurons within UCP were organized into vertical queues, with adjacent neurons expressing DSCAM and positioned in close proximity. Notably, these queue-like structures were not observed in the *Dscam^-/-^* brain. Together with our live-imaging studies, these observations suggest that DSCAM-DSCAM interactions between neighboring neurons contribute to synchronized radial migration by limiting forward progression of trailing neurons. This mechanism is consistent with a role for DSCAM in regulating inter-neuronal spacing and preserving sequential order within migrating neuronal cohorts.

### DSCAM preserves sequential order within radial queues

Based on the *Dscam* expression, as revealed by ISH, and its known role in regulating cell-cell spacing and axon guidance ^30,31^, we speculated that DSCAM-mediated interactions maintain the radial queue of migrating neurons. We therefore predicted that when a fast-migrating neuron from the LCP approaches a stalled neuron at the base of UCP, its radial migration would slow, preventing it from overtaking. To test this, we performed time-lapse imaging focused on the base of UCP, ∼150 µm below the CP/ML border. Newly-arriving WT neurons migrated quickly from LCP (magenta arrowheads, Fig. 2a WT 1:00-4:00) and approached a neuron stalled at the base of UCP (green arrowheads, Fig. 2a WT, 5:00-8:00), and then halted migration (Fig. 2a, WT 8:00-12:00). This is in marked contrast to *Dscam^-/-^* neocortex where the fast-migrating neuron (magenta arrowheads) bypassed stalled neurons (green arrowheads) at the base of the UCP (Fig. 2a, *Dscam^-/-^* 1:00-3:30). Quantification revealed that in WT brains, most trailing neurons (85.9±7.6%) halted when reaching the suspended leading neurons (Fig. 2b) and localized in close range of -7.0±6.1 µm relative to the leading neurons in WT (Fig. 2c). In *Dscam^-/-^* cortex, only 27.6±24.0% of trailing neurons halted, and most of them bypassed their leading neurons, with an average position of 9.6±13.7 µm ahead of the leading neurons during the imaging (Fig. 2b and 2c). A diagram illustrates that DSCAM is required for leading neurons to restrict the migration of their following neurons, forming queues of migrating neurons (Fig. 2d). These data indicate that the queue may be the radial assembly of migrating neurons.

**Fig. 2.**
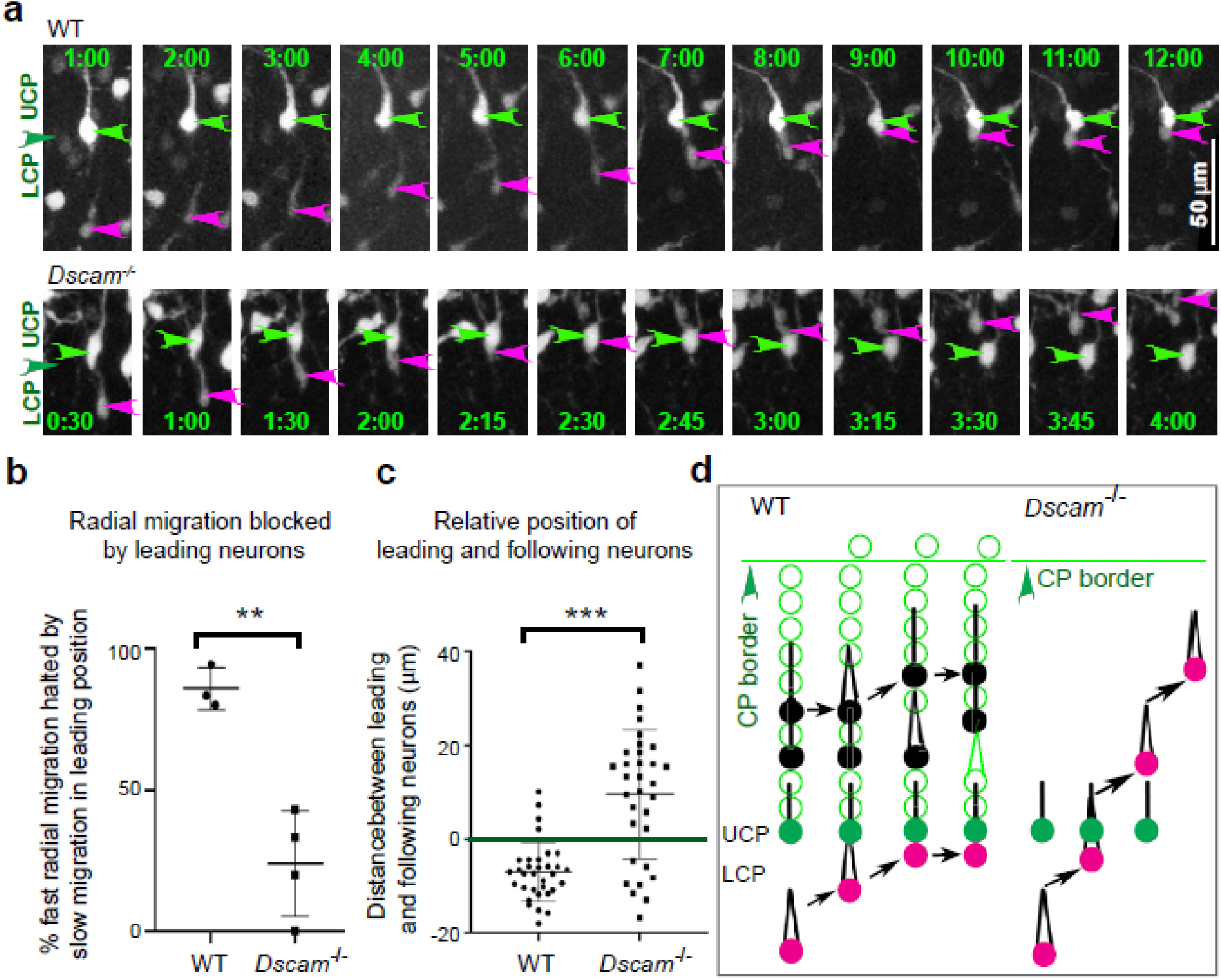
Stalled radial migration of neurons in a leading position directly impedes the fast radial migration of trailing neurons. **a** Time-lapse images of electroporated neurons in WT cortex (top) and *Dscam^-/-^* cortex (bottom). The fast-migrating trailing neurons are labeled with a magenta arrowhead. The leading neurons are labeled with a green arrowhead. **b** Quantification of migration behavior. Y-axis, the % of fast-migrating neurons blocked by the leading neurons. Data points represent individual animals. Student’s t-test. **c** Quantification of relative position of leading and trailing neurons after being encountered. The leading neuron position is considered 0 and is labeled by a green line. **p<0.01, and ***p<0.001. Data points represent pairs of a leading and trailing neuron from 4 WT and 4 *Dscam^-/-^* brains. Mann-Whitney U test. **d** Diagram showing that stalled neuron in the leading position (green) at the UCP bottom halts the approaching fast-migrating trailing neuron (magenta) to maintain a migratory queue in the wild-type cortex, whereas the leading neuron (green) is passed by trailing neuron (magenta) in the *Dscam^-/-^* cortex.

### DSCAM is required in both leading and trailing neurons to maintain migratory order

DSCAM can function as a homophilic cell adhesion molecule^30^. In some developmental contexts, *trans* interactions between DSCAM molecules on neighboring neurons can antagonize Cadherin-mediated adhesion and promote neuronal self-avoidance ^34^. In addition, DSCAM can also interact with several heterologous ligands^36^. It therefore remains unclear whether maintaining the migratory order depends on *Dscam* expression in leading or trailing neurons, or in both. To address this question, we employed inducible knockout (iKO) of *Dscam* by introducing Cre and GFP-expression plasmids into *Dscam^fl/fl^* embryos via IUE at E15. GFP^+^ neurons distributed broadly in *Dscam^fl/+^* cortex, while GFP^+^ iKO neurons accumulated in a narrower region near the CP/ML border in *Dscam^fl/fl^*cortex, similar to what we observed in the *Dscam^-/-^* brain (Fig. 3a and 3d). These results indicate that loss of *Dscam* in trailing neurons prevents their migration from being stalled by leading WT neurons, suggesting a cell-autonomous requirement in following neurons. As a control, this phenotype of *Dscam* iKO neurons can be rescued by co-expression of full-length *Dscam* (Fig. 3b and 3d). Notably, a truncated form of DSCAM lacking the intracellular domain (DSCAM ΔICD) also rescued the phenotype of *Dscam* iKO neurons (Fig. 3c and 3d). This shows that the DSCAM ectodomain is sufficient for queue formation, the radial assembly of migrating neurons.

**Fig. 3.**
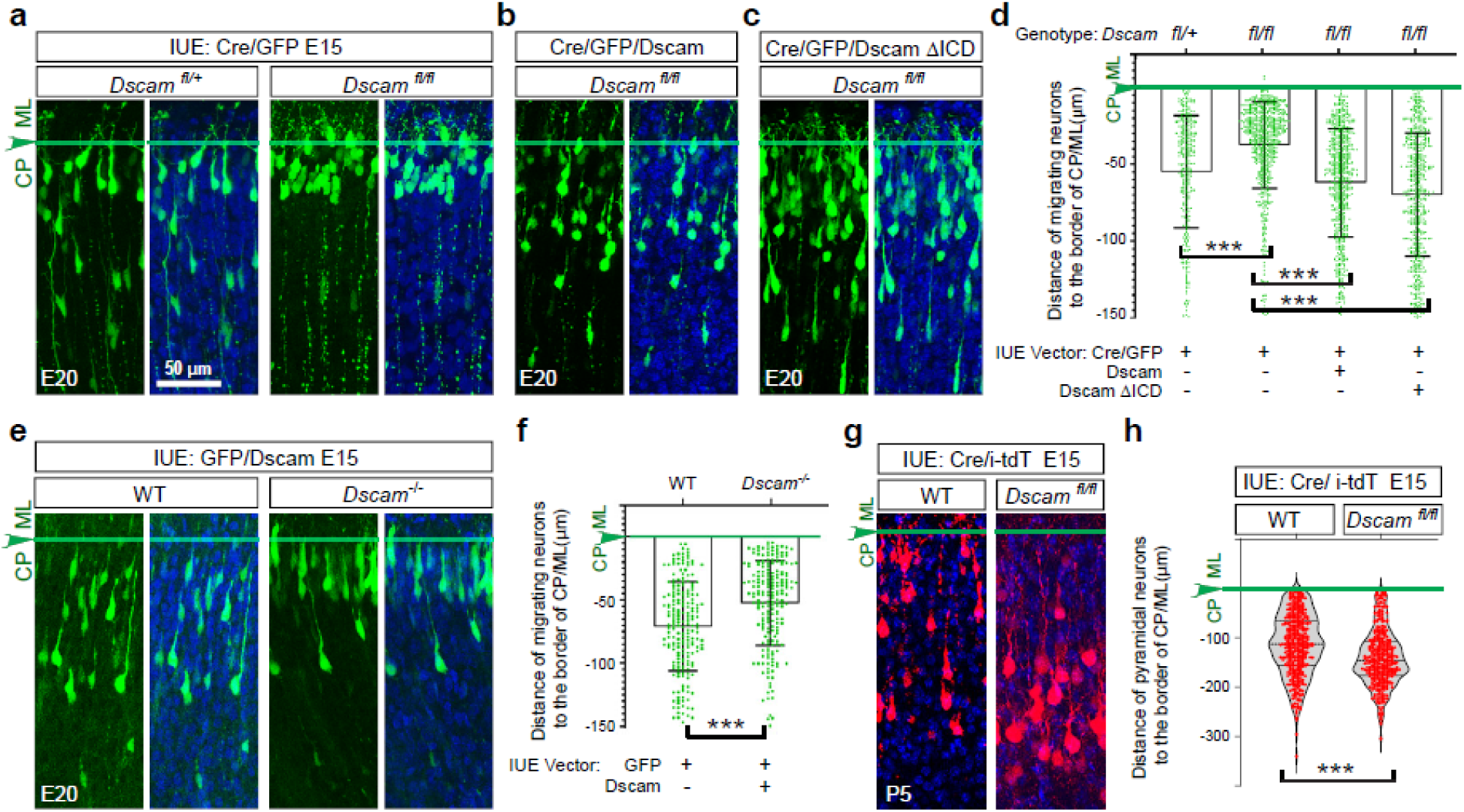
Cell-autonomous function of DSCAM in radially migrating cortical neurons. **a** Neocortical sections of E20 embryos. The distribution of GFP^+^ neurons (green) and DAPI-labeled nuclei (blue) following IUE of Cre and GFP expression vectors into *Dscam^fl/+^* and *Dscam^fl/fl^* embryos at E15. **b,c** Neocortical sections following re-introduction of full-length DSCAM (**b**) or DSCAM ΔICD (intracellular domain was deleted, **c**) in *Dscam^fl/fl^*;*Cre* neurons. **d** Quantification of the distance of GFP^+^ neurons to the CP/ML border in 3 *Dscam^fl/+^*, 4 *Dscam^fl/fl^*, 4 full-length DSCAM, and 4 DSCAM ΔICD brains. Kruskal-Wallis test. **e** Distribution of GFP^+^ neurons in WT or *Dscam^-/-^* cortical plate with DAPI counterstaining following IUE of *Dscam* overexpression at E15. **f** Quantification of the distance of GFP^+^ neurons to the CP/ML border in 3 WT and 4 *Dscam^-/-^* brains. Mann-Whitney U test. **g** Distribution of RFP^+^ neurons (Red) in P5 UCP with DAPI counterstaining (blue) following IUE of Cre and Cre-inducible RFP vectors in *Dscam^fl/fl^*mice at E15. **h** Quantification of the distance of WT neurons (6 brains) and *Dscam^-/-^*neurons (6 brains) to the CP/ML border. Mann-Whitney U test. ns, not significant, and ***p<0.001. Scale bar: 50 µm. The arrowhead and green line point to the CP/ML border.

We next tested whether *Dscam* expression in leading neurons is also required to constrain trailing-neuron migration. A *Dscam*-expression plasmid was introduced into control or *Dscam^-/-^*neural progenitors by IUE. In control cortex, GFP^+^/DSCAM^+^ neurons were broadly distributed, whereas in *Dscam^-/-^* cortex, these neurons accumulated near the CP/ML border (Fig. 3e and 3f). This indicates that *Dscam^-/-^*leading neurons cannot block DSCAM^+^ trailing neurons. Together with the iKO results (Fig. 3a and 3d), these data support a model in which DSCAM is required in both the leading and trailing neurons to maintain orderly radial migration.

Finally, we examined long-term consequences of *Dscam* loss in neocortical neuronal positioning. Cre and a Cre-inducible RFP reporter were introduced into *Dscam^fl/fl^* embryos via IUE at E15, and cortical organization was examined at postnatal day 5 (P5), when most pyramidal neurons had completed radial migration and settled in the cerebral cortex. Compared with WT controls, RFP^+^ (*Dscam* iKO) neurons were positioned significantly deeper within *Dscam^fl/fl^* cerebral cortex (Fig. 3g and 3h), indicating impaired progression to the superficial layers.

### DSCAM suppresses F-actin assembly and cadherin-mediated adhesion

Time-lapse imaging revealed that as the leading neuron moves towards the CP/ML border and creates space, the trailing neuron promptly extends its leading process, thickening its proximal end to form the PCDLP ^13^ (Fig.1d, upper panel 1:00-2:00, green brackets). The extension of PCDLP is necessary to trigger the saltatory nucleokinesis of radial migration (Fig. 1d, upper panel 2:00-4:00). These observations suggest that the positioning of leading neurons suppresses PCDLP expansion, thereby limiting nucleokinesis. To validate this, we assayed the PCDLP of migratory neurons in the UCP and LCP (Fig. 4a). We found that 14.7±7.5% of migrating neurons in the WT UCP form a PCDLP, significantly lower than 94.6±3.1% in *Dscam^-/-^* UCP, 82.8±8.4% in WT LCP, and 92.6±5.9% in *Dscam^-/-^* LCP (Fig. 4b). Most migrating neurons in the WT UCP lack a PCDLP, supporting the notion that leading neurons may restrict radial migration of following neurons by inhibiting PCDLP expansion in following neurons.

**Fig. 4.**
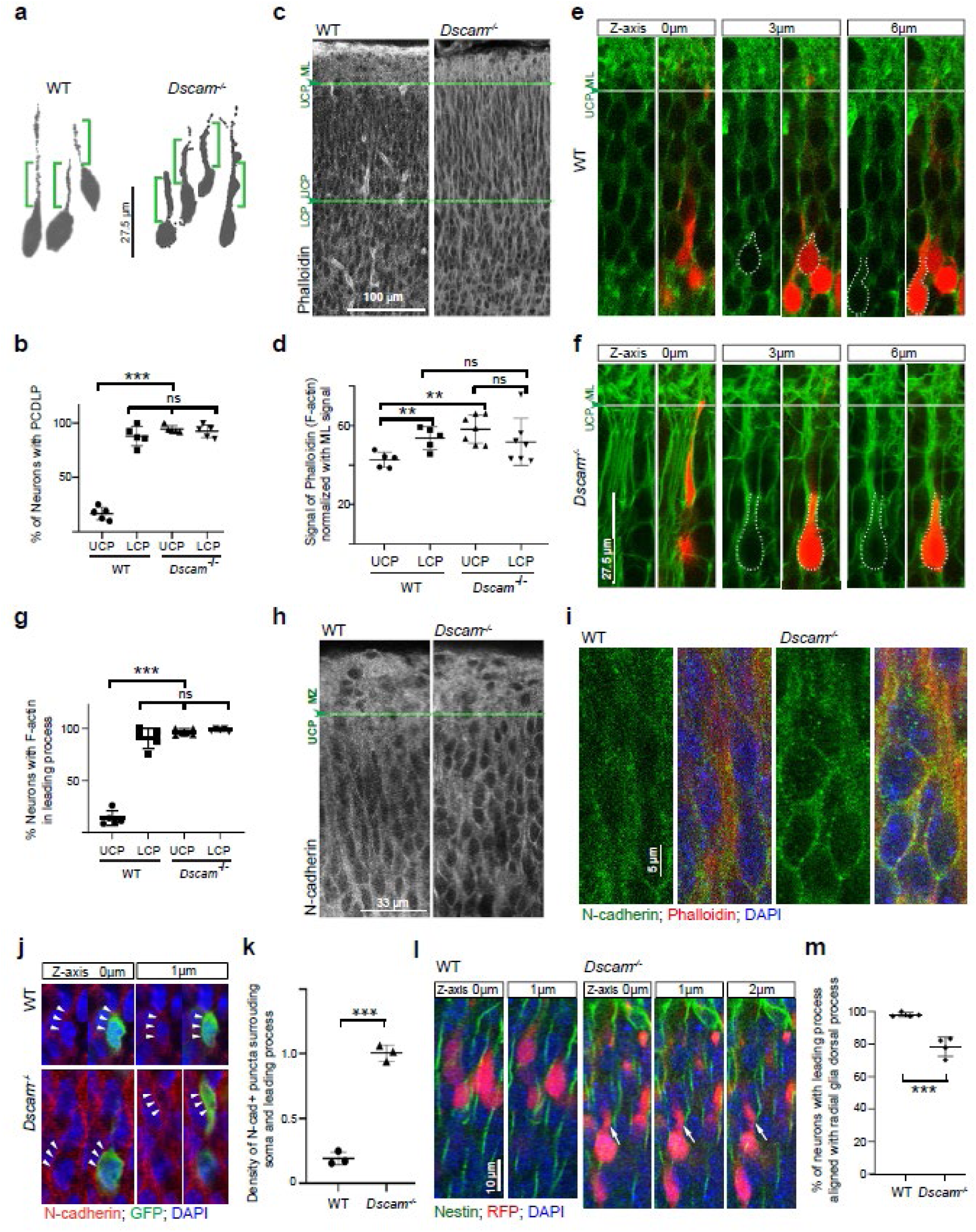
DSCAM influences F-actin assembly and N-cadherin adhesion puncta in the UCP. **a** tracing the radial-migrating neurons in WT and Dscam^-/-^ cortices. Brackets label the proximal end of the leading process. **b** Quantification of the % of migrating neurons with PCDLP (leading process thickness ≥1.2 µm measured at 5-10 µm ahead of soma) in the UCP. Ordinary one-way ANOVA. **c** Sections through the cortical plate of E18.5 WT and *Dscam^-/-^* embryos, stained with phalloidin for F-actin. **d** Quantification of F-actin signal intensity in the UCP and LCP of both genotypes, normalized to staining of the molecular layer (ML). Ordinary one-way ANOVA. **e,f** Higher magnification of neocortex showing F-actin in individual WT (**e**) and *Dscam^-/-^*neurons (**f**) in the UCP. Z-axis labels the z-positions of confocal imaging relative to the original layer (0µm). **g** Quantification of the % of migrating neurons with F-actin in PCDLP or cell body. Ordinary one-way ANOVA. **h** Neocortical sections showing N-cadherin staining in E19.5 CP. **i** Higher magnification of N-cadherin staining (Green) in the E19.5 UCP. **j** N-cadherin surrounding migrating neuron at E19.5 in the WT and *Dscam^-/-^* UCP. Arrowheads label the edge of the leading process and soma. **k** Quantification of N-cadherin puncta density (/µm) surrounding neuronal soma and leading process. Student’s t-test. **l** Relative position of migrating neurons and the dorsal process of radial glial cells (Nestin staining, green) in E19.5 UCP. Z-axis labels the z-positions of confocal imaging relative to the original layer (0µm). Arrows label the leading process not along the radial glia. **m** Quantification of the % of leading processes attached to the dorsal process of radial glial cells. Data points represent individual mice. Student’s t-test. ns, not significant, **p<0.01, and ***p<0.001.

Because F-actin assembly leads to the protrusion of lamellipodia and PCDLP expansion^13^, we next examined F-actin using phalloidin staining in E18.5 cortex. F-actin intensity in the UCP was normalized to the ML F-actin intensity, which remained consistent across all samples (Fig. 4c). F-actin levels in WT UCP were significantly lower than those in *Dscam^-/-^*UCP, while the F-actin signal is comparable in the WT LCP, *Dscam^-/-^*UCP, and *Dscam^-/-^* LCP (Fig. 4c and 4d). High-magnification images revealed that in contrast to low level of F-actin in radial-migrating neurons (Fig. 4e), F-actin appeared to be highly assembled throughout soma and leading process in *Dscam^-/-^*neurons in UCP (Fig. 4f). Notably, the percentage of radial-migrating neurons in the WT UCP exhibiting F-actin signal in leading process was remarkedly lower than that in the WT LCP, *Dscam^-/-^* UCP and *Dscam^-/-^* LCP (Fig. 4g). The low F-actin level is consistent with fewer lamellipodia in the proximal end of leading process, which suggests the expansion of the PCDLP and nucleokinesis was hindered in WT UCP.

Cadherin-mediated adhesion is known to promote and stabilize F-actin assembly during lamellipodia protrusion^37,38^. Previous studies have shown that DSCAM can antagonize Cadherin adhesion by Cdh2^16^, Cdh3, or Cdh6^34^ during neuronal patterning. Hence, DSCAM is thought to function by “masking” cadherin adhesion. N-cadherin (Cdh2) is essential for proper radial migration^39^, as its adhesion around neurons promotes F-actin assembly beneath the adhesion sites^37^. This makes N-cadherin an ideal candidate for studying the potential mechanistic link between DSCAM and cadherin adhesion. Immunohistochemistry showed that N-cadherin localization was diffuse and broad in the WT UCP, while strong and well-defined puncta were observed in the *Dscam^-/-^* UCP (Fig. 4h and 4i). This is consistent with increased Cadherin-mediated adhesion in the absence of DSCAM^16^.

Additional images revealed that N-cadherin puncta (arrowheads pointed puncta, Fig. 4j) were present on soma (Fig. 4j, *Dscam^-/-^*, 0µm) and leading process (Fig. 4j, *Dscam^-/-^*, 1µm), potentially forming adhesions with other neurons ( Fig. 4i and 4j), with 1.01±0.06 puncta per µm on the edge of soma and leading process. In WT, N-cadherin puncta density was significantly lower (Fig. 4j and 4k). These results support a model wherein DSCAM is required to block Cadherin-mediated adhesion among migrating neurons. In this model, without N-cadherin adhesion, F-actin disassembles in the following neuron to dismantle its nucleokinesis in the WT UCP.

During cortical development, nascent neurons undergo radial migration along radial glia. Previous studies have shown that *Dscam* is highly expressed in migrating immature neurons but largely absent in radial glial cells, as determined by single-cell RNA sequencing (Suppl. Fig. 2^40^). This differential expression permits Cadherin-mediated adhesion between radial glia and migrating neurons, thereby allowing radial glia to function as a migration scaffold. Since some nascent neurons formed cadherin adhesion with neighboring neurons in the *Dscam^-/-^*cortex (Fig. 4i and 4j), we reasoned those nascent neurons might not form adhesion with radial glia and could migrate independently of radial glial dorsal process. Indeed, radial glia labeled by anti-nestin staining showed a significantly higher percentage of neurons with leading processes not aligned with radial glia dorsal process in the *Dscam^-/-^* UCP (Fig. 4l and 4m).

### DSCAM generates intercellular clefts that limit cadherin-mediated adhesion

To address the question of how DSCAM might block Cadherin-mediated adhesion in more detail, we first transfected HeLa cells (lack of endogenous *Dscam*) with N-cadherin-BFP, either alone or together with DSCAM-mNeonGreen (Fig. 5a). Without DSCAM, neighboring HeLa cells formed N-cadherin adhesive puncta (Fig. 5a). In the presence of DSCAM, surface N-cadherin was separated by a narrow cleft with an average width of 1.36±0.56 µm between cells (Fig. 5a). N-cadherin localizes to the cell surface, indicating that DSCAM neither excludes N-cadherin from the cell surface nor affects the extension of N-cadherin-containing protrusions into the cleft. Instead, DSCAM may block N-cadherin processes from encountering each other across the cleft. Indeed, the cleft between two N-cadherin^+^ HeLa cells was filled with thin mNeonGreen^+^ DSCAM strands (Fig.5a). These features indicate that DSCAM fibers between two cells may interact to prevent N-cadherin processes from making physical contact. It is also possible that dense DSCAM fibers may inhibit N-cadherin from adhesion *in trans*.

**Fig. 5.**
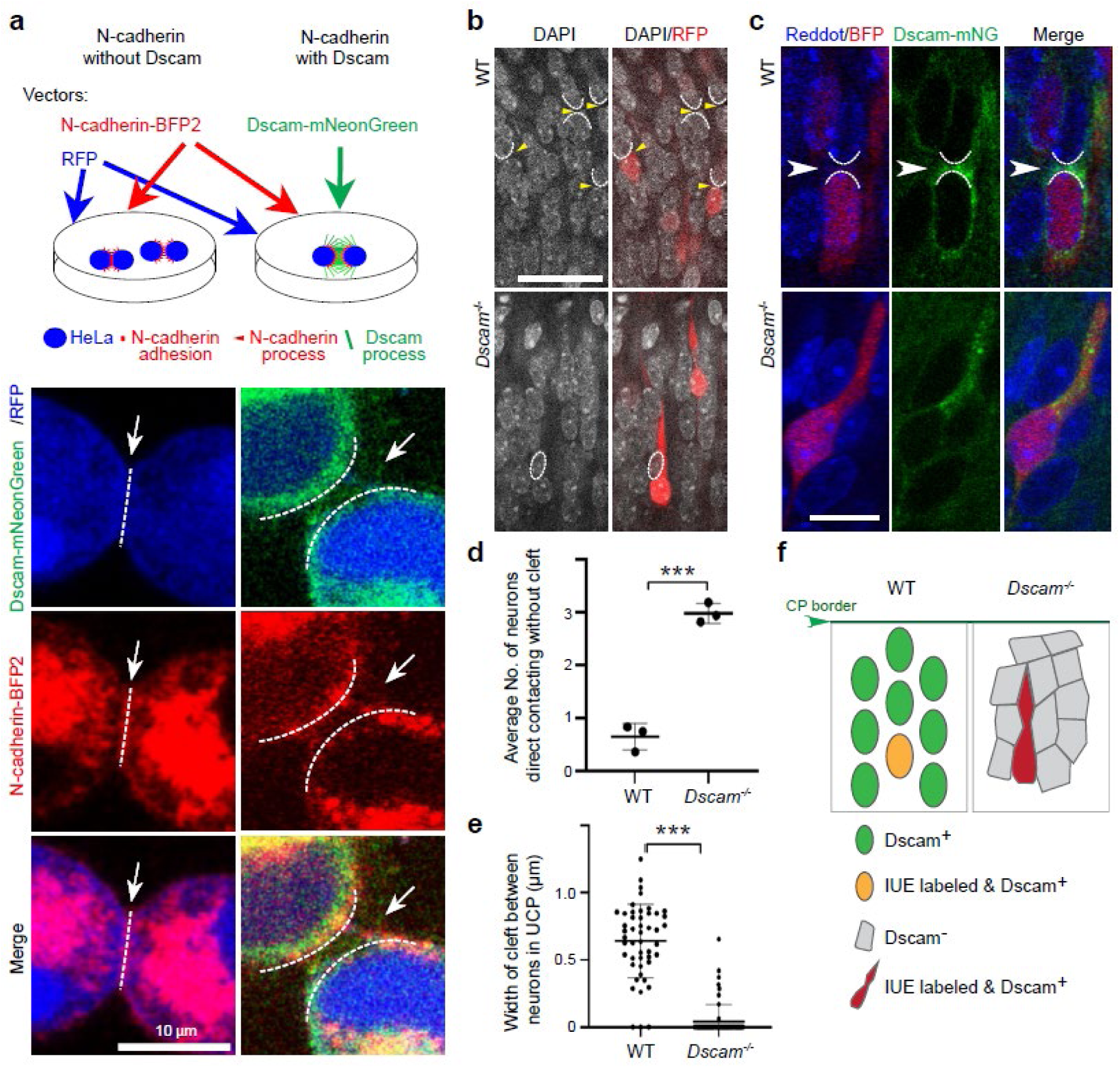
DSCAM is required to keep neurons separated. **a** Diagram showing the transfection of N-cadherin-BFP, with or without DSCAM-mNeonGreen into HeLa cells. In the absence of DSCAM (lower left panel), a white dashed line and arrow labeled N-cadherin (red) puncta form between cells. In the presence of DSCAM protein (green; lower right panel), the white dashed lines and arrows label the narrow cleft between HeLa cells to show the position of N-cadherin (red). DSCAM-positive fibers (green) localized to the gap between cells. Scale bar, 10µm. **b** Neocortical sections showing WT neurons separated by gaps (indicated by yellow arrowheads) in the E19.5 UCP, *Dscam^-/-^* neurons packed with diminished gaps. RFP labels radial-migrating neurons by IUE. Scale bar, 30µm. **c** Localization of DSCAM (green) in radial-migrating neurons (red) in the E19.5 UCP with Reddot counterstaining (blue) following IUE of BFP (Red) and DSCAM-mNeonGreen (Green) at E14.5. Arrowhead points to the DSCAM (green) in the gap between neurons. White dashed lines label the edges of the neurons. Scale bar, 10µm. **d** Quantification of an average number of neurons directly in contact with other migrating neurons with diminished gaps. Student’s t-test. Data points represent individual animals. **e** Quantification of the average width of gaps between the IUE-labeled migratory neurons and their adjacent neurons. Data points represent individual neurons from 3 WT mice and 3 *Dscam^-/-^* mice. Mann-Whitney U test. ***p<0.001. **f** Diagram demonstrates a model wherein DSCAM functions to keep neurons separated in the UCP. In *Dscam^-/-^* cortical plate, neurons are packed with diminished gaps. Interestingly, *Dscam* expression in a single neuron via IUE cannot form gaps that separate it from adjacent Dscam^-^ neurons.

We next asked whether DSCAM also generates gaps between migrating neurons. In the WT UCP, neurons were isolated from each other (Fig. 5b, Suppl. Fig. 3). Each neuron directly contacted less than one adjacent neuron (Fig.5d). The width of clefts separating adjacent neurons was 0.64±0.27µm (Fig.5e). In contrast, *Dscam*^-/-^ neurons were squeezed more closely together (Fig. 5b, Suppl. Fig. 3), each neuron directly contacted ∼3 other neurons (Fig.5d), and there was little space of ∼0.04±0.12µm between neurons (Fig.5e). Does this suggest that DSCAM separates radial-migrating neurons from each other?

To answer this question, DSCAM-mNeonGreen was expressed in migratory neurons via IUE to visualize DSCAM *in vivo*. We found that mNeonGreen^+^ DSCAM covered the soma (Red), filling the cell-cell clefts surrounding the soma in WT (Fig. 5c). Similar as observed in HeLa cells, DSCAM staining forms dense and thin fibers that grow from the WT cell surface (Fig. 5c). Interestingly, in *Dscam^-/-^* brain, mNeonGreen^+^ DSCAM remained inside migrating neurons (Red), which directly contacted the surrounding *Dscam^-/-^* neurons (Fig 5c). Together, these results indicate that DSCAM is required for generating clefts between neurons to separate them (Fig. 5b-5f).

### DSCAM physically blocks cadherin-mediated adhesion among radial-migrating neurons

To test if DSCAM dominantly suppresses cadherin-mediated adhesion among migrating neurons, we overexpressed N-cadherin-eGFP via IUE at E14.5 and assayed the impact of extra N-cadherin adhesion on neuron clustering in the E18.5 CP. In WT brains, radial-migrating neurons expressing N-cadherin-eGFP did not form a cluster in the UCP, and only 11±6% of migratory neurons contacted the other migratory neurons (Suppl. Fig. 4a and 4b). In contrast, migratory neurons with N-cadherin-eGFP formed clusters (>3 cells), with approximately 53±5% of migratory neurons clustered in *Dscam^-/-^*UCP (Suppl. Fig. 4a and 4b). These results indicate that DSCAM physically suppresses cadherin-mediated adhesion in the UCP, and that overexpression of N-cadherin cannot overcome the DSCAM-mediated adhesion block.

### Balanced DSCAM and UNC5c function regulates radial migration

A previous study showed that DSCAM directly binds to UNC5c via extracellular interactions^41^. To assess UNC5c’s role in radial migration, a previously validated *UNC5c*-specific shRNA^41^, was introduced into E14.5 neural progenitors via IUE. Neurons with *UNC5c* knockdown exhibited delayed migration across UCP at E19 (Fig. 6a), and the average distribution of migrating neurons across the UCP was -65.4±35.3µm below the CP/ML border, compared to - 47.8±32.1µm in controls (Fig. 6b). In contrast, UNC5c overexpression (OE) via IUE caused over-migration, with neurons accumulating below the CP/ML border, with an average distribution of -32.7±27.0µm. These results indicate that UNC5c promotes radial migration across the UCP.

**Fig. 6.**
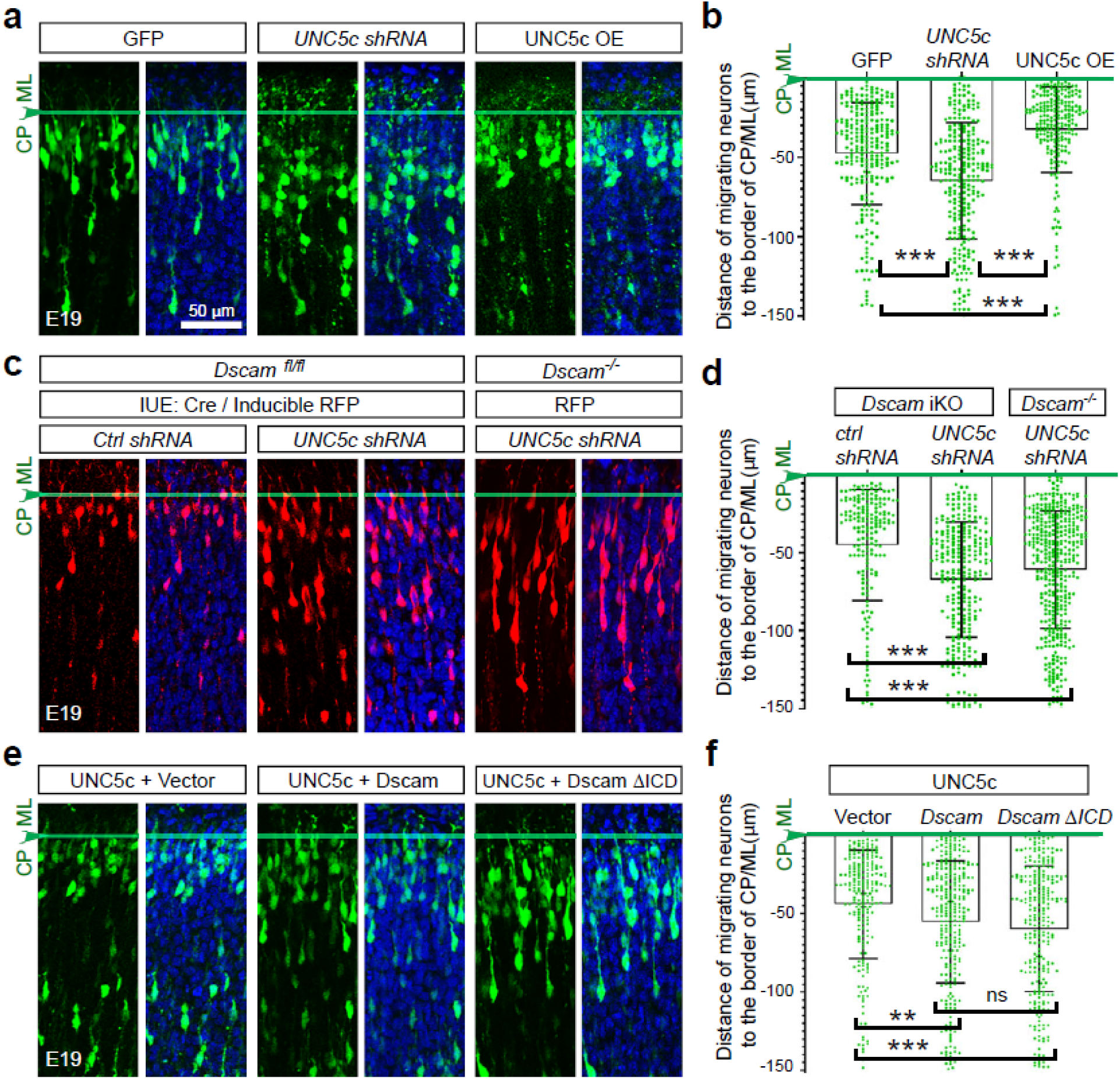
DSCAM and UNC5c cooperate to regulate radial migration in the UCP. **a** Neocortical sections showing distribution of GFP+ neurons in the E19 UCP with DAPI counterstaining (blue) following IUE of Control *shRNA*, *UNC5c shRNA*, or UNC5c overexpression (OE) vectors, respectively, with GFP vector at E14.5. The results were repeated in 3 brains for each condition. **b** Quantification of the distance of GFP^+^ neurons to the CP/ML border. **c** Distribution of RFP^+^ neurons in the E19 UCP with DAPI counterstaining following IUE of inducible RFP and Cre vectors with Control *shRNA* or *UNC5c shRNA* into *Dscam^fl/fl^*, and *UNC5c shRNA* into *Dscam^-/-^*brains at E14.5. Control shRNA was tested in 3 brains. UNC5c shRNA was tested in 4 *Dscam^fl/fl^* brains and 4 *Dscam^-/-^* brains. **d** Quantification of distance of RFP^+^ neurons to the CP/ML border. **e** Distribution of GFP^+^ neurons in E19 UCP with DAPI counterstaining (blue) following IUE of UNC5c overexpression vector with empty vector, WT *Dscam*, or *Dscam-ΔICD* vector at E14.5, respectively. The results were repeated in 3 brains for each condition. **f** Quantification of the distance of GFP^+^ neurons to the CP/ML border. Kruskal-Wallis test. ns, not significant, **p<0.01, and ***p<0.001.

Since both DSCAM and UNC5c regulate radial migration in the UCP, it remains unclear how they coordinate this process. We next tested whether DSCAM contributes to the delayed radial migration induced by *UNC5c* knockdown. At E14.5, *UNC5c shRNA* was co-transfected with Cre and inducible RFP into *Dscam^fl/fl^* NPCs via IUE, resulting in labeled neurons being *Dscam* iKO and *UNC5c* knockdown. Distribution of these neurons in the UCP was markedly broader than that in *Dscam* iKO neurons alone (Fig. 6c and 6d). These results suggest that UNC5c promotes migration independently of DSCAM, as the UNC5c phenotype persists in the absence of DSCAM.

Since UNC5c OE caused neurons’ over-migration to the dorsal border of the CP (Fig. 6a and 6b), we next asked whether increasing DSCAM expression could counteract this over-migration phenotype. UNC5c and DSCAM OE constructs were co-transfected at E14.5. Double OE of either UNC5c and DSCAM, or UNC5c and DSCAM ΔICD rescued over-migration caused by UNC5c OE (Fig. 6e and 6f). These results indicate a functional balance between UNC5c-mediated repulsion and DSCAM-mediated restraint of migrating neurons in UCP.

## DISCUSSION

Radial migration of cortical neurons is essential for establishing the characteristic “inside-out” development of cerebral cortex. Previous studies have primarily focused on the terminal step of migration at the cortical plate/marginal layer (CP/ML) border, where migrating neurons detach from radial glial fibers and complete migration^15,16^. While this process is critical for laminar positioning, the mechanism by which the sequential order of migrating neurons is preserved during migration remains unclear. In this study, we report that neurons form and maintain ordered queues and that DSCAM organizes migrating neurons into the radial queues within UCP, thereby preserving their sequential order and contributing to inside-out corticogenesis.

### Radial assembly of migrating neurons forms a queue to maintain the neuronal order

Here, we show that radial migration in the UCP is distinct from that in the LCP. Before entering UCP, neurons migrate at a constant and rapid rate. In contrast, migration within the UCP becomes markedly reduced and intermittent, with neurons undergoing episodic saltatory nucleokinesis interspersed with prolonged pauses. Time-lapse imaging further revealed that leading neurons block trailing neurons from expanding the proximal cytoplasmic dilation of the leading process, thereby impeding nucleokinesis in the UCP. In this way, leading neurons temporarily restrain trailing neurons, synchronizing radial migration in the UCP. DSCAM is essential for the difference in migratory behavior, as *Dscam^-/-^* neurons fail to exhibit this characteristic slowing behavior and instead continue migrating rapidly to the CP/ML boundary. These migratory behaviors, together with the observation that radial-migrating neurons form evenly spaced vertical queues spanning the UCP, suggest that DSCAM maintains inter-neuronal spacing and organizes neurons into migratory queues.

*Dscam* ISH revealed the overall architecture of the queues, in which individual DSCAM^+^ neurons are organized into vertical columns across the UCP. In contrast, DSCAM^+^ neurons in LCP are isolated, with little or no contact between neighboring DSCAM^+^ cells. Using *in-utero* electroporation, we labeled radial-migrating neurons and found that these neurons were DSCAM^+^ and positioned either within the vertical DSCAM column in the UCP or as isolated signal in the LCP. The distinct migratory behaviors and corresponding *Dscam* expression patterns in the UCP and LCP suggest that the vertical DSCAM column maintains synchronized migration and the sequential order of neurons. Accordingly, we propose that leading neurons transiently impede trailing neurons via DSCAM-mediated interactions, thereby forming a radial assembly that preserves neuronal synchronization and migratory order.

If the proposed model is correct, it is reasonable to predict that fast-migrating neurons ascending through the LCP should be halted upon reaching the UCP at the base of the vertical column. Moreover, if a neuron were suspended at the base of UCP, it should be possible to observe this neuron blocking the fast-approaching neuron. Time-lapse imaging confirmed these predictions, as neurons approached the base of the UCP, the rapid migration ceased. Fast-migrating neurons approaching from the LCP were immediately halted beneath suspended preceding neurons at the bottom of the UCP. These observations confirmed that the vertical DSCAM column represents a radial assembly of migrating neurons. Within the column, leading neurons restrain trailing neurons to synchronize their movement. Newly arriving neurons are blocked at the base of the vertical column, where they join and extend the assembly.

### The neuronal sequential order in the queue is converted into the “inside-out” cortical morphogenesis

Our previous studies suggest that radial migration is not simply a process of neuronal translocation, but an ordered transitional state that converts temporal sequential order into cortical lamination^16^. During migration, neurons maintain a fixed order within radial queues. However, upon reaching the CP/ML boundary and completing migration, later-arriving neurons bypass earlier post-migratory neurons and enter the emerging cortical layer in ML. This local reversal of relative position transforms the initially linear migratory order into the characteristic “inside-out” architecture of the cerebral cortex^16^. Thus, the radial queue of migrating neurons serves as a critical intermediate structure that preserves and transfers neuronal order during corticogenesis.

To directly examine the role of neuronal DSCAM in corticogenesis, we disrupted DSCAM function in migrating neurons by introducing Cre into *Dscam^fl/fl^*neurons using IUE. As expected, DSCAM deficiency in either the leading or trailing neuron prevents the formation of organized migratory queues. Together, these gain- and loss-of-function studies indicate that DSCAM functions cell-autonomously in migrating neurons, where its expression is required in both the leading and trailing neurons to maintain orderly radial migration. Without this queue-based constraint, neurons prematurely reached the CP/ML border and positioned in the deeper layers. Thus, in the absence of DSCAM-mediated order, newly arriving neurons no longer preserve their relative sequence during migration and instead intermingle with earlier-arrived neurons. A similar phenotype is observed in *Dscam^2j/2j^* cortex, where accelerated arrival of upper-layer neurons intermingle at the CP/ML border, leading to thinning of upper cortical layers^16^. Together, these findings support a model in which a radial queue serves as a temporal ordering mechanism that preserves the sequential progression of migrating neurons, thereby enabling the “inside-out” organization at the CP/ML boundary.

### DSCAM-mediated intercellular clefts block Cadherin-mediated adhesion

Our results show that DSCAM from adjacent neurons forms a narrow cleft that blocks N-cadherin adhesion, thereby maintaining neuronal spacing. The narrow cleft was first found between two DSCAM^+^/N-cadherin^+^ cells, rather than adhesive puncta, when DSCAM was not expressed. DSCAM^+^ fibers are long and thin, extending from opposing cellular cortices to fill this cleft. N-cadherin^+^ fibers extend from both sides, flanking the clefts, but do not connect to each other. This arrangement suggests that DSCAM does not exclude N-cadherin from the cleft interface but instead blocks direct N-cadherin-mediated contacts.

Similar intercellular gaps are present between migrating neurons in the queue, where cadherin-mediated adhesion is sparse. DSCAM fiber extended from soma separates the neurons from each other. In contrast, DSCAM-deficient neurons exhibit excessive neuronal adhesion and fail to maintain spacing. Moreover, DSCAM is required from both sides. Radial glial cells express little, if any, DSCAM, allowing migrating neurons to maintain N-cadherin-mediated adhesions along them as a migration scaffold. Lastly, DSCAM^+^ fiber *in trans* interactions does not promote tight cell-cell adhesions or F-actin assembly. So, DSCAM-mediated interactions inhibit PCDLP expansion and subsequent nucleokinesis.

The rapid response of migrating neurons within the queue also supports a physical blockade model. When leading neurons migrate and create space, trailing neurons immediately occupy it (Fig. 1d). While, as trailing neurons encounter the base of suspended leading neurons, their nucleokinesis abruptly halts (Fig. 2a, WT). These rapid responses are difficult to explain solely through a gradual signaling mechanism and instead suggest direct physical interference between adjacent neurons.

If the DSCAM-mediated cleft functions as a physical blockade, it should not be rescued by increasing N-cadherin level. Consistent with this prediction, N-cadherin overexpression in WT brains fails to overcome the DSCAM-mediated cleft, neither enhancing cadherin-mediated adhesion nor promoting neuronal clustering. Previous studies similarly show that, although N-cadherin is required for radial migration, its overexpression does not drive excessive migration of nascent pyramidal neurons^39^. In contrast, in *Dscam^-/-^*brains, N-cadherin overexpression elevates neuronal adhesion, leading to neuronal clustering. In the absence of DSCAM, higher N-cadherin levels directly strengthen cell-cell adhesion. Together, these findings support the idea that DSCAM-mediated clefts function as a physical barrier that blocks N-cadherin-mediated neuronal adhesion, thereby maintaining the spacing of migrating neurons.

### DSCAM counterbalances UNC5c to regulate radial migration in UCP

DSCAM-mediated clefts likely function as a physical barrier; however, the physical blockade alone may not explain DSCAM’s dose-dependent functions on axon terminal size, synapse formation, etc^42^. DSCAM may therefore regulate cortical thickness through additional gradual or dose-dependent mechanisms beyond its role as a physical barrier that maintains intercellular spacing.

In addition to DSCAM-mediated spacing, we identify UNC5c as a key regulator of neuronal movement within the UCP. UNC5c, a Netrin-1 receptor, binds DSCAM through its extracellular domain^41^. We find that UNC5c drives migrating neurons toward the CP/ML border in a dosage-dependent manner. Knockdown of UNC5c disperses neurons throughout the UCP in both WT and DSCAM-deficient cortices, indicating that UNC5c-mediated repulsion acts independently of DSCAM. Functionally, UNC5c provides driving force for upward migration, whereas DSCAM constrains the progression of trailing neurons to preserve migratory order. Thus, formation of a DSCAM-dependent migratory queue requires UNC5c-driven displacement towards the boundary. In contrast to physically blocking N-cadherin, DSCAM expression level counterbalances UNC5c to fine-tune radial migration in UCP in a dosage-dependent manner.

The distinct effects of DSCAM on UNC5c and N-cadherin may reflect fundamental differences between cadherin-mediated adhesion and receptor-ligand signaling. Cadherin-mediated adhesion requires cadherins to be present on both adjacent cell membranes, making it highly sensitive to intercellular spacing. In contrast, receptor-ligand signaling only requires receptors on the responding cell, with ligands diffusing through the extracellular space. Therefore, the DSCAM-mediated cleft may effectively prevent cadherin binding between neighboring membranes without interfering with the diffusion of ligands such as Netrin-1.

Together with previous studies on Reelin signaling at the CP/ML border, our findings support a model in which multiple mechanisms cooperate to establish inside-out cortical architecture. Reelin reduces adhesion at the CP/ML boundary, ensuring that neurons terminate migration only at the appropriate superficial location. DSCAM preserves the sequential order of migrating neurons during transit through the UCP, while UNC5c promotes their upward displacement. Through the coordinated actions of these pathways, the temporal order of neuronal migration is converted into the layered organization of the cerebral cortex.

In summary, our study identifies that the radial assemblies of migrating neurons form queues spanning the UCP as a previously unrecognized intermediate structure in the morphogenesis of upper cortical layers. Our study establishes that DSCAM generates clefts between neurons, blocking the cadherin adhesion, to maintain spacing and order during radial migration. These findings provide new insight into how collective neuronal migration contributes to cortical lamination and may help elucidate the pathological mechanisms underlying neurodevelopmental disorders.

## MATERIALS AND METHODS

### Mice and genotyping

In this study, we used *Dscam^2j/2j^* mutant mice^33^ for loss-of-function studies and time-lapse imaging. *Dscam^2j/2j^* mutant mice do not make a protein product of *Dscam*. The *Dscam^2j^* mutation contains a 4-base-pair duplication in exon 19 of *Dscam*, leading to a frameshift during translation. *Dscam^2j/2j^* mutant mice cannot give birth to offspring. To overcome the breeding barrier, *Dscam^2j/+^* mice (hereon referred to as *Dscam^+/-^*) were bred to generate wild-type (*Dscam^+/+^*), heterozygotes (*Dscam^+/-^*), and mutant (*Dscam^-/-^*) mice. The *nestin-CreERT2* mouse expresses tamoxifen-induced Cre recombinase under the *nestin* promoter in the neural progenitor cells (NPCs) during cortical development. The *Ai14* mouse expresses tdTomato, a red fluorescent protein, in a Cre-dependent manner. *Dscam^+/-^*mice were bred into *nestin-CreERT2* and *Ai14* mice to generate *Dscam^-/-^/nestin-CreERT2/CAG-tdTomato* progeny with tamoxifen-inducible *nestin*-promoter-restricted tdTomato expression in NPCs. tdTomato expression in NPCs was induced by a dose of tamoxifen injection into dams bearing E14.5 embryos. To determine whether Dscam has the cell-type autonomous function for maintaining the radial assembly of migrating neurons, *Dscam^flox^* (*Dscam^fl/+^*) mice (Stock No. 17689^43^) were bred to generate heterozygotes (*Dscam^fl/+^*) and homozygotes (*Dscam^fl/f/^*) offspring. Thus, *cre* overexpression plasmid was transfected into NPCs via *in-utero* electroporation at E14/15, thereby achieving inducible *Dscam* knockout in heterozygous (*Dscam ^fl/+^; cre*) or homozygous (*Dscam ^fl/fl^; cre*) pyramidal neurons.

Genomic DNA from embryos was purified for genotyping. Embryonic tail tissues were digested in 300 µL TES buffer (50 mM Tris pH 8.0, 0.5% SDS, 0.1 M EDTA, in 1X PBS) with proteinase K at 37 °C overnight, followed by addition of 200 µL 5 M NaCl. Samples were centrifuged at 12,000 rpm to remove the precipitates. 200 µL pure isopropanol was added to 300 µL supernatant to precipitate DNA. DNA pellets were washed with 500 µL 75% ethanol before being dissolved in 20 µL ultra-pure water. DNA surrounding the *Dscam^2j^*mutation was amplified by PCR (forward primer: 5’-gccctgtggtatttgctggtgtg-3’; reverse primer: 5’-Gatgggcaaatgtcaaaggtcaaa-3’). PCR products of *Dscam^2j^* offspring were sequenced with either primer to determine the presence of the 4-base-pair insertion. For *Dscam^flox^*genotyping, primers (forward primer:5’-agcaaaagcaccatgattgacag-3’ and reverse primer: 5’-caactgacttaatgagtggag-3’) were used.

All animal handling was in accordance with the policy and guidance of the Institutional Animal Care & Use Committee (IACUC) at the University of Michigan, as well as the protocol approved by the IACUC. All mice were in a dedicated temperature-controlled (20°C) animal vivarium with a 12-hour light/dark cycle.

### Plasmids

To make the N-cadherin-EGFP expressible in radial-migrating neurons, N-cadherin-EGFP was excised with NheI (blunt end) and NotI (sticky end) from pCMV-N-cadherin-EGFP (Addgene, plasmid#18870) and subsequently inserted into the PB-CAG plasmid to generate PB-CAG-N-cadherin-EGFP. For N-cadherin fused with 2XTagBFP2, EGFP in PB-CAG-N-cadherin-EGFP was replaced by 2XTagBFP2, which was excised from pCMV-2XTagBFP2 with SmaI and NotI, to generate PB-CAG-N-cadherin-2XTagBFP2. To create a DSCAM and mNeonGreen fusion protein, *Dscam* was excised from pCMV-mouse-*Dscam* (Addgene, plasmid#18737) with EcoRI and XhoI and then inserted into pCS2-mNeonGreen plasmid. To overexpress *Dscam*-mNeonGreen in migrating neurons, *Dscam*-mNeonGreen was excised from pCS2-*Dscam*-mNeonGreen using EcoRI and NotI and inserted into the PB-CAG vector, creating pCAG-*Dscam*-mNeonGreen. To generate truncated DSCAM lacking its intracellular domain, the intracellular domain was removed with AFLII and Smal digestion to produce pCAG-*Dscam*-ΔICD. pTT3-UNC5c-flag (Addgene, plasmid#72196) was used for UNC5c overexpression. UNC5c shRNA vector was a kind gift from Dr. Guofa Liu.

### *In-utero* electroporation (IUE) of mouse embryos

IUE was performed as previously described^13,44^. The pregnant dam mouse (E14.5) was anesthetized with isoflurane, and an incision was made to expose the embryos in the uterus. Plasmid solution (∼1μg/μl) containing 0.01% w/v Fast Green dye was injected into one of the lateral ventricles with a glass micropipette. Each embryo head in the uterus was placed between the leads of a Tweezertrode Electrode system (5 mm in diameter, BTX, Harvard Bioscience) and administered five electrical pulses (50 V, 50 ms duration, 950 ms intervals) using an electroporator (ECM830, Harvard Bioscience) to transfect the dorsal neural progenitor cells (NPCs). The embryos were then returned to the abdominal cavity. The muscles and skin were sutured, respectively. The pregnant dam was administered carprofen (5 mg/kg, i.p.) for pain management and was monitored till revived.

### Live-cell time-lapse confocal microscopy of brain sections

Brain slice culture and time-lapse confocal imaging were carefully described in our 2023 Bio-protocol paper^45^. The time-lapse confocal imaging datasets were also used in our previous paper (PMID: 35672151), which focused on the stop of radial migration outside of the CP/ML border. Briefly, embryonic brains were coronally sectioned into 300 μm slices in ice-cold artificial cerebrospinal fluid using a vibratome (Leica Microsystems, Wetzlar, Germany). Brain slices were then mounted in ice-cold Matrigel and incubated at 37°C for 30 minutes to solidify. Then, the brain slice was continuously perfused with DMEM bubbled with a gas mix (95% O_2_, 5% CO_2_) at 37°C on the microscope stage. Time-lapse imaging was performed using the Leica TCS SP5 confocal microscope with an HC Fluotar L 25x/0.95 W VISIR immersion objective lens at 5-minute intervals. After the time-lapse imaging, the embryos were genotyped as described above.

### Fluorescence staining and immunohistochemistry of brain sections

Brains were fixed with 4% paraformaldehyde in 1xPBS overnight. Then, brains were kept in 30% sucrose overnight, then embedded in optimal cutting temperature (O.C.T.) media (Tissue-Tek^TM^, Sakura), and frozen at -80 °C. Brains were sectioned coronally at 50 µm thickness, permeabilized with 0.2% Triton X-100 in 1xPBS, blocked with 2% normal donkey serum in 1xPBS with 0.2% Triton X-100, and incubated overnight at 4 °C with primary antibody or phalloidin in 1xPBS containing 2% normal donkey serum and 0.2% Triton X-100. The antibodies and dyes used are: (i) mouse anti-N-cadherin antibody (1:100, GC-4 clone, Sigma, C3865), (ii) goat anti-Nestin (1:200, R&D, AF2736), and (iii) Phalloidin-647 (1:500, Abcam, Ab176759). Sections were rinsed and probed with secondary antibodies (1:500, Jackson ImmunoResearch). The brain sections were then counterstained with DAPI or NucRed Live 647 (1:1000, Invitrogen, R37106), rinsed, and mounted in 50% glycerol before visualization. Images were captured using a Leica SP5 confocal microscope.

### *In-situ* hybridization (ISH) on E19 brain sections

*Dscam* mRNA ISH was performed using published probe sequences. Probe sequences were amplified from *Dscam* cDNA [catalog no. 18737, Addgene] and subcloned into a pTOPO vector. DIG-labeled cRNA probes were generated with a DIG RNA labeling kit (Roche, Basel, Switzerland), as previously described ^16^. DIG-labeled cRNA probes were produced by *in vitro* transcription from the *Dscam* cDNA template. ISH was performed on fresh-frozen mouse brain sections mounted on microscope slides, which were fixed and permeabilized with proteinase K. The sections were then incubated with DIG-labeled cRNA probes, rinsed, and incubated with the anti-DIG-AP (alkaline phosphatase) antibody (Roche). The signal was detected using an AP substrate (NBT/BCIP; Roche).

### HeLa cell culture and transfection

HeLa cells were cultured in DMEM with 10% FBS in a humidified cell culture hood at 37 °C and 5% CO_2_. For transfection and imaging, HeLa cells were cultured on poly-D-lysine-coated coverslips. N-cadherin-GFP [1.5 µg plasmid; catalog no. 18870, Addgene] or N-cadherin-2XTag-BFP2 (1.5 µg plasmid)/DSCAM-mNeonGreen (1.5 µg plasmid) were transfected, respectively, with Lipofectamine 3000 (Invitrogen, Thermo Fisher Scientific). Transfected HeLa cells were incubated in a humidified cell culture hood at 37 °C and 5% CO_2_ for 2 days. Then, the HeLa cells were fixed on the coverslip with 4% paraformaldehyde, counterstained with NucRed Live 647 (1:1000, Invitrogen, R37106), rinsed, and mounted on a glass slide for imaging under the Leica SP5 confocal microscope.

### Statistical Analysis

GraphPad Prism 11 was used for data analysis. Numbers were presented as the means ± standard deviation of means. The investigator quantifying the images was blinded to genotypes or plasmid transfection. Normality was assessed using the Shapiro-Wilk test. Parametric datasets were analyzed using the original one-way ANOVA followed by Student’s t-test or directly using Student’s t-test. Nonparametric datasets were analyzed using the Kruskal-Wallis test followed by Dunn’s multiple-comparisons test or the Mann-Whitney U test. P< 0.05 was considered statistically significant.

## Supporting information

supplmental informatino

## Acknowledgments

The authors thank Dr. Jack Parent, Dr. Guofa Liu, and Dr. Jun Wu for sharing reagents. This study was supported by the NIH R01MH119346 to Roman Giger.

## Author contributions

T.Y. conceived and designed the study; T.Y., M.W.V., E.J.C., T.H., and X-F.Z. performed the experiments; T.Y., W-C.C., Y.W., P.G.F., R.J.G. and X-F.Z. analyzed the data; T.Y. and X-F.Z. wrote and revised the manuscript with input from all authors.

## Competing interests

The authors declare no competing interests.

## Additional information

Supplementary information is available online.

## Notes

### Competing Interest Statement

The authors have declared no competing interest.

### Summary of Updates

The major new results incorporated into the revised manuscript are summarized below. 1. We now show the Dscam expression pattern in the lower cortical plate using in situ hybridization (Suppl Fig 1d). 2. We used a truncated Dscam construct lacking the cytosolic domain to rescue the distribution defects of migrating neurons caused by Dscam knockout (new Figure 3c and d). 3. We analyzed postnatal day 5 WT and Dscamfl/fl brain samples to show the cortical layering defects caused by Dscam KO (new Figure 3g and h). 4. We now show the N-cadherin puncta surround the proximal cytoplasmic dilation of the leading process and the cell body of the migrating neurons (Fig 4j). 5. We used a Nestin antibody to label dorsal radial glial fibers (new Figure 4l and m), demonstrating that loss of Dscam causes radial-migrating neurons to deviate from radial glial fibers. 6. We used IUE of green and red fluorophores to differentially label neurons, demonstrating DSCAM-dependent spacing between adjacent neurons (Suppl Fig 3). 7. We studied UNC5c functions in radial migration in the UCP and its relation to DSCAM (new Figure 6).

