## Supplementary material for "DSCAM preserves neuronal order during radial migration by antagonizing N-Cadherin adhesion and UNC5c repulsion": supplmental informatino

**The file includes:**

Suppl Fig 1 to 4,

Suppl Refences.

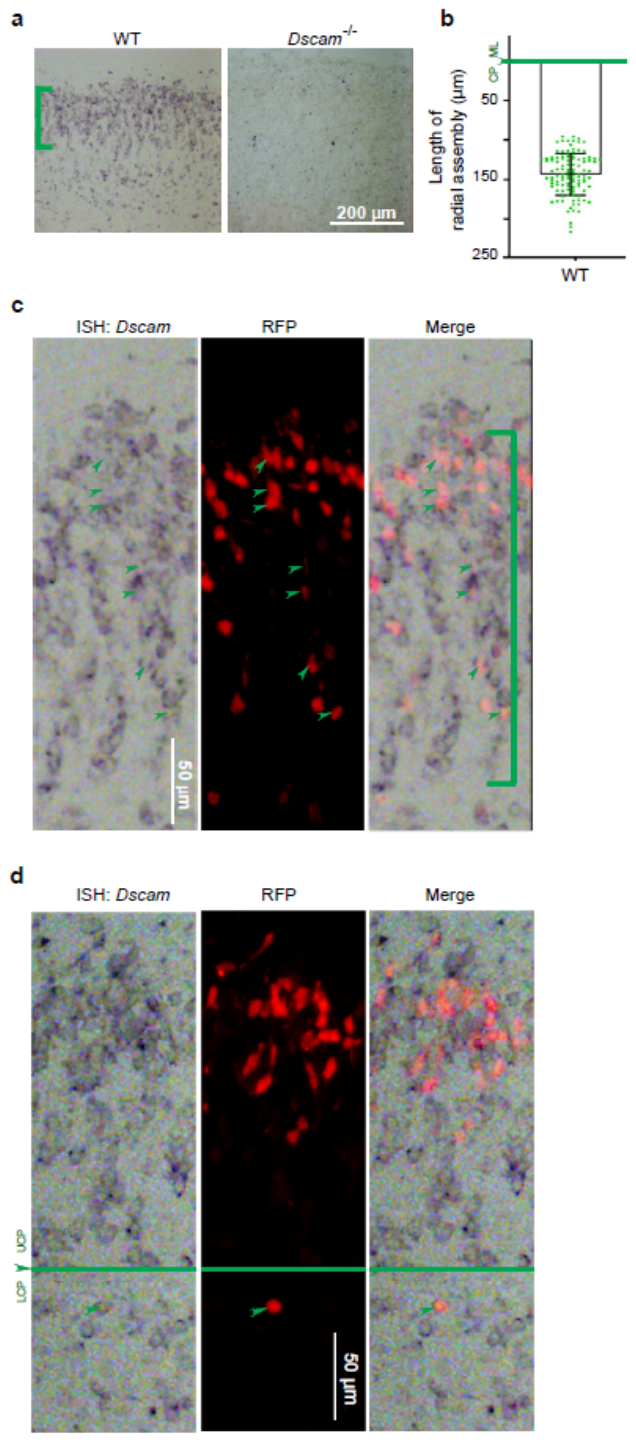

**Suppl Fig. 1**, ***in-situ* hybridization detects *Dscam* mRNA-positive queues near the CP/ML border. a** Cortical sections of E19 WT and *DSCAM^-/-^* mice stained for *Dscam* transcripts by in situ hybridization. **b** Quantification of the length of the vertical column of *Dscam* in cortical plates is shown by green brackets labeled in (**a** and **c**). Data points represent individual queues of radial assemblies in 3 brains. **c** Enlarged images show that WT neurons (red), labeled using IUE at E14.5, form queues perpendicular to the border of the dorsal cortical plate and express *Dscam* (green arrowheads). **d** WT neuron with *Dscam* mRNA signal in LCP. Green arrowheads label in-situ hybridization signal, RFP^+^ neurons, and merged images via IUE at E14.5.

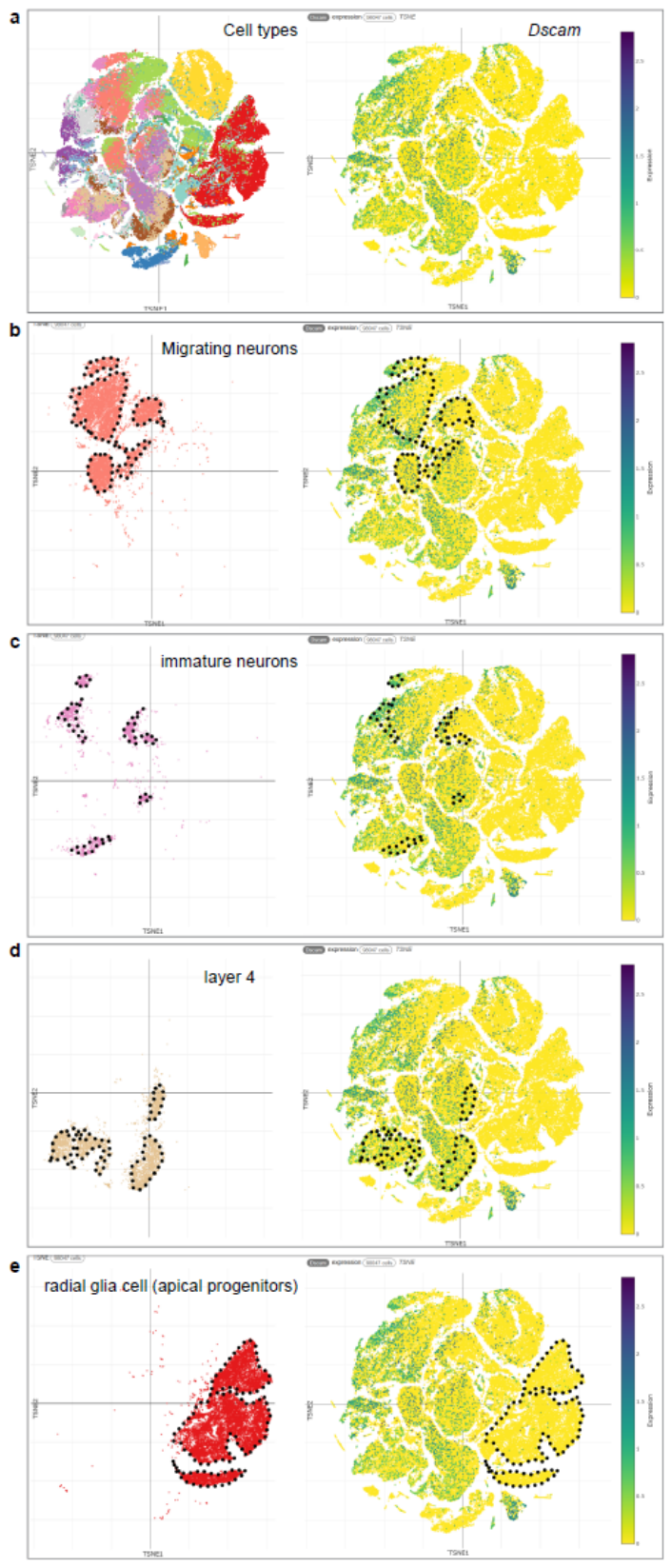

**Suppl Fig. 2,** ***Dscam* expression during mouse cortex development.** TSNE plot of integrated snRNAseq datasets generated from E10, E11, E12, E13, E14, E15, E16, E17, E18 (S1), E18 (S3), P1, and P4 mouse cerebral cortex. Color coding is used to show cell types and expression level. **a** Color-coded cell types (left) and distribution of *Dscam* expression (right). *Dscam* expression in migrating neurons (**b**), immature neurons (**c**), and layer 4 neurons (**d**). Very little *Dscam* expression was detected in radial glia cells (**e**). Datasets were originally described in^1^.

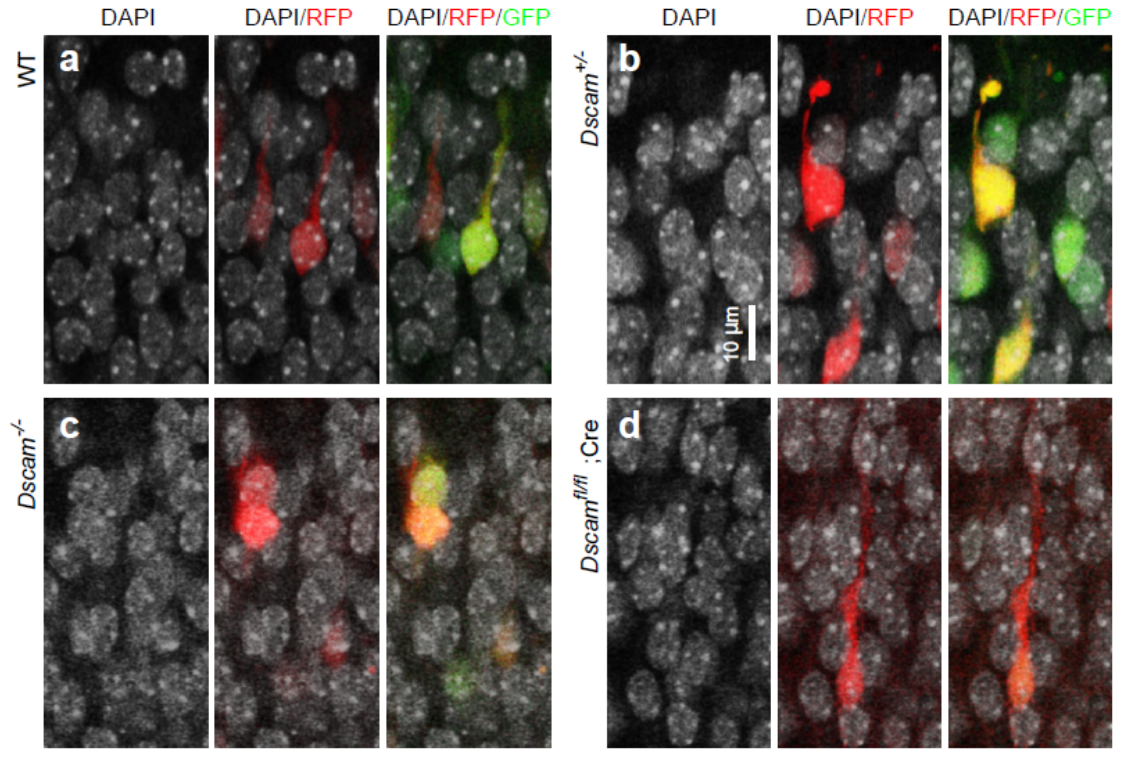

**Suppl Fig. 3, DSCAM is necessary to keep neurons separated.** IUE was performed at E14.5 to label neurons with GFP and/or RFP. The migrating neurons (red or green) on E19 neocortical sections are separated from the other neurons by narrow gaps in WT (a) and *Dscam^+/-^* genotypes (b). The gaps disappear in the *Dscam^-/-^* mutant (c). Cre-induced conditional *Dscam* knockout neuron (d), labeled by Cre-induced RFP, directly contacts the other neurons in an otherwise WT cortical plate.

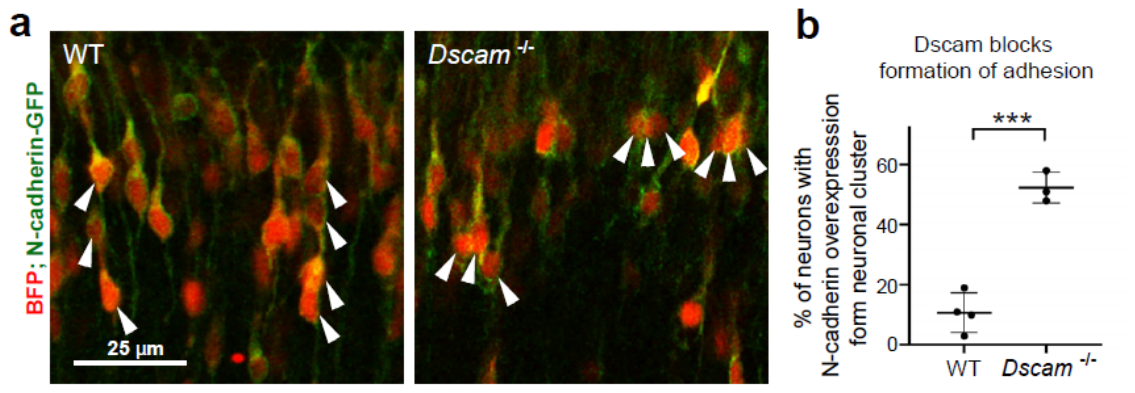

**Suppl Fig. 4, DSCAM blocks N-cadherin adhesion among radial-migrating neurons. a** Coronal brain section, on which BFP and N-cadherin-GFP were overexpressed in WT and *Dscam^-/-^* brains via IUE at E15. At E19, the clustering of RFP^+^ migrating neurons with N-cadherin-GFP in WT and *Dscam^-/-^* UCP was compared. **b** Quantification of % of clustered migrating neurons (≥3 neurons) with N-cadherin-GFP overexpression in UCP. Data points represent individual mice. Student’s *t-test*, ***p<0.001.
